# CENP-A chromatin prevents replication stress at centromeres to avoid structural aneuploidy

**DOI:** 10.1101/2020.09.01.277103

**Authors:** Simona Giunta, Solène Hervé, Ryan R. White, Therese Wilhelm, Marie Dumont, Andrea Scelfo, Riccardo Gamba, Cheng Kit Wong, Giulia Rancati, Agata Smogorzewska, Hironori Funabiki, Daniele Fachinetti

## Abstract

Chromosome segregation relies on centromeres, yet their repetitive DNA is often prone to aberrant rearrangements under pathological conditions. Factors that maintain centromere integrity to prevent centromere-associated chromosome translocations are unknown. Here, we demonstrate the importance of the centromere-specific histone H3 variant CENP-A in safeguarding DNA replication of alpha-satellite repeats to prevent structural aneuploidy. Rapid removal of CENP-A in S-phase, but not other cell cycle stages, caused accumulation of R-loops with increased centromeric transcripts, and interfered with replication fork progression. Replication without CENP-A causes recombination at alpha-satellites in an R-loop-dependent manner, unfinished replication and anaphase bridges. In turn, chromosome breakage and translocations arise specifically at centromeric regions. Our findings provide insights into how specialized centromeric chromatin maintains the integrity of transcribed noncoding repetitive DNA during S-phase.

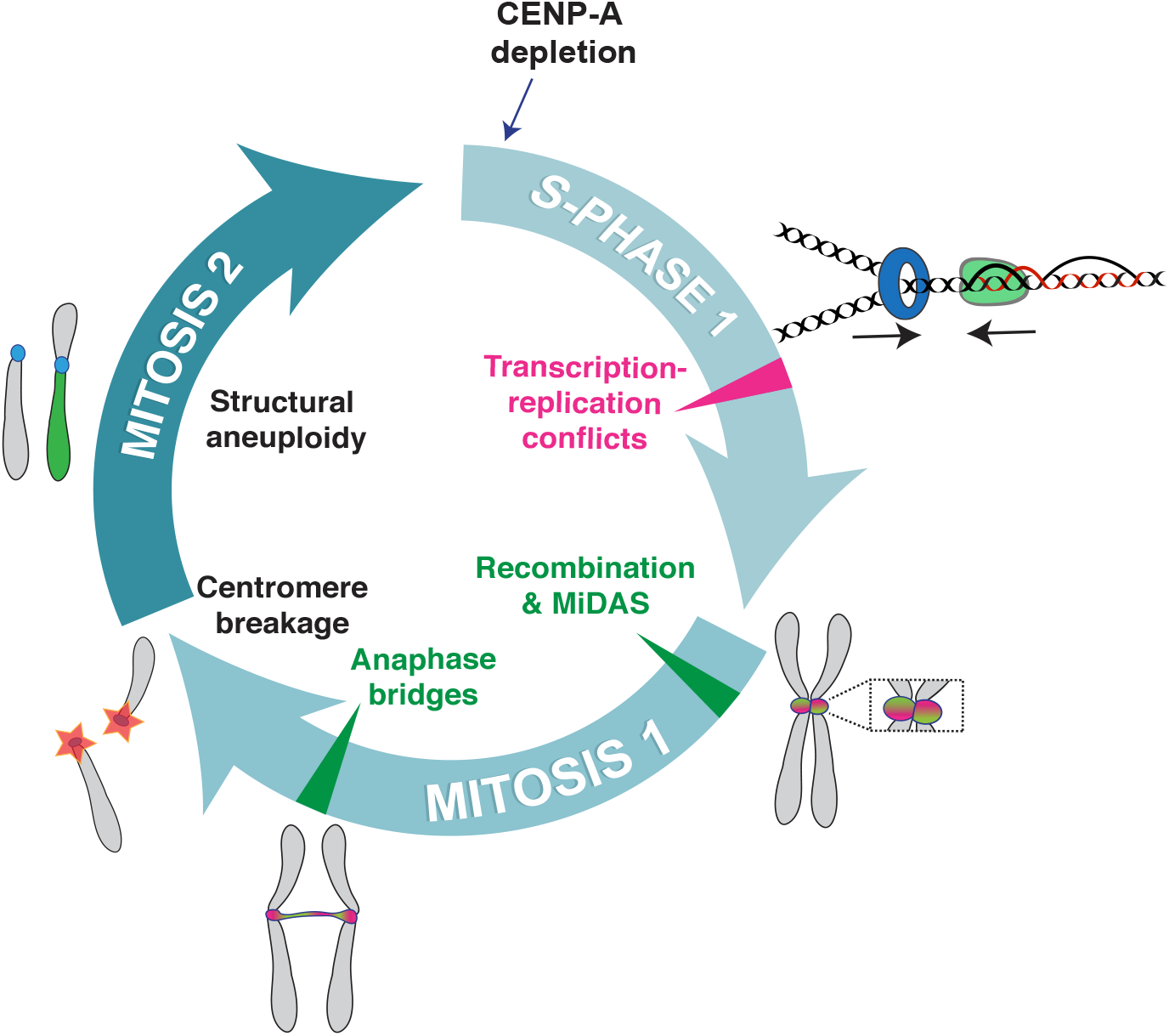

## Introduction

Centromeres hold a paradox: their essential function to support accurate chromosome segregation is evolutionarily conserved yet their DNA sequence is rapidly evolving (1–3). Human centromeres span ~0.3-5 Mb with repetitive alpha-satellite DNA arranged in tandem and then reiterated as homogenous higher order repeat (HOR) blocks (4). Evolution of centromere repeats has been modelled by short and long-range recombination events such as gene conversion, break-induced replication, crossing over and other mutagenic processes (2, 3, 5, 6). Changes in centromere size may contribute to meiotic drive (2), but may also elicit genome instability (7) by increasing the rate of chromosome mis-segregation (8). While recombination at centromeres is suppressed in meiosis (9), it is present in somatic cells and enhanced in cancerous cells (10) where whole-arm chromosome translocations are prevalent (11, 12). The mechanism(s) that prevent(s) breakage and translocations at these highly repetitive centromere sequences remain elusive.

Human centromeres are epigenetically specified by the histone H3 variant CENP-A (13, 14), which acts as a locus-specifying seed to assemble kinetochores for mitotic functions (15). While pericentromeric regions form heterochromatin, the core centromere harbors euchromatic characteristics with active transcription throughout the cell cycle, where long non-coding RNAs act *in cis* to contribute to centromere functions (16, 17). To date, it is unknown how a) transcription (17), b) recombination (10), c) late replication (18, 19) and d) propensity to form non-B-DNA and secondary structures (20–22), all features commonly associated with human fragile sites (23), are regulated to maintain the integrity of centromeric repeats. Here we identified a critical role for CENP-A in the maintenance of centromeric DNA repeats by repressing R-loop formation during DNA replication. Replication stress caused by rapid CENP-A depletion in S-phase leads to delayed centromeric mitotic replication and formation of chromosome bridges, which trigger overall genome instability through chromosome breakage and translocations in the subsequent cell cycle. We demonstrated that in addition to enabling mitosis, CENP-A maintains chromosome integrity by facilitating replication of centromere repeats.

## Results

We previously demonstrated that long-term CENP-A loss promotes alpha-satellite recombination events (10), suggesting a potential role for CENP-A in the maintenance of the integrity of centromere repeats, beyond its role in kinetochore formation and spindle stability (14, 24). As telomeres and rDNA possess specialized mechanisms that prevent their repetitive sequences from instability in S-phase (25, 26), we hypothesized that CENP-A plays a role in suppressing alpha-satellite fragility during DNA replication. To test this, we used a system that allows rapid removal of endogenous CENP-A-containing nucleosomes using an Auxin Inducible Degron (AID) (Figure 1A and S1A) in hTERT-immortalized, non-transformed, diploid Retinal Pigment Epithelial (hTERT-RPE-1) cells (27). We first monitored consequences of CENP-A removal on nucleosome stability using stably expressed SNAP-tagged H4. As previously reported (28), when SNAP-H4 is labeled by TMR in G1, the majority of signals accumulate at centromeres (Figure 1B, C), since CENP-A nucleosomes assemble during late M/early G1 (29), whereas canonical nucleosomes assembly is coupled to DNA replication. Auxin (IAA) addition led to the disassembly of CENP-A-containing nucleosomes as shown by rapid loss of previously incorporated centromeric SNAP-tagged H4 concomitant to ^AID^CENP-A (Figure 1B-D). However, new incorporation of H4 at centromeres at any phases of the cell cycle was not altered by CENP-A removal in G1 (Figure S1B, C), as assessed by H4^SNAP^ quenching/release and H4K5Ac CUT&RUN-qPCR (30), indicating that nucleosomes dynamics were not affected by acute CENP-A depletion. Considering that a large majority of each human centromere is occupied with H3 nucleosomes with ~200 interspersed CENP-A nucleosomes (31), we expect the impact of short-term CENP-A depletion on the integrity of centromeric chromatin to be minimal.

**Figure 1.**
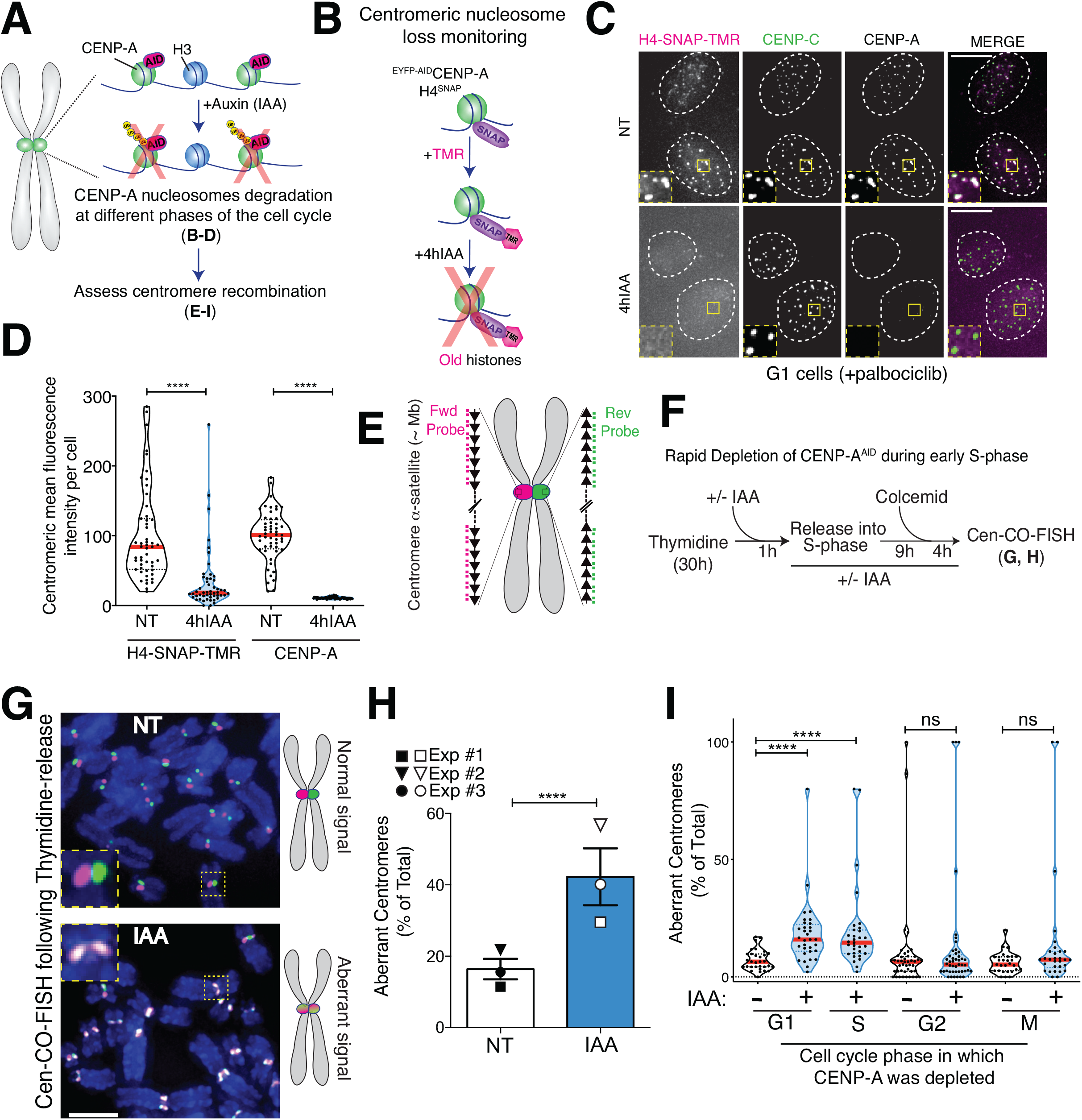
Rapid removal of CENP-A during S-phase causes centromeric nucleosomes loss and increases centromere recombination. (A) Schematic of the inducible degradation system to deplete endogenous CENP-A after addition of Auxin (IAA) in RPE-1 cells. (B) Strategy to monitor nucleosome loss by identifying the H4^SNAP^ labeled with the TMR fluorophore prior to the induction of CENP-A depletion. (C) Representative immunofluorescence images of ^EYFP-AID^CENP-A + H4^SNAP^ G1-arrested cells showing centromeric nucleosome loss after 4 h IAA. Scale bar: 10 μm (D) Quantification of CENP-A and TMR (previous deposited histone H4 pool) mean fluorescence intensity at centromeres after 4 h IAA per cell. n>50 cells per condition. Red lines represent the median. Mann-Whitney test: **** p<0.0001. (E) Schematic illustration of the Cen-CO-FISH centromeric DNA probes. Hybridization by unidirectional PNA probes differentially labels the forward or reverse strands of each sister chromatid. Each black arrow symbolizes a HOR in the alpha-satellite array. (F) Synchronization strategy for the Cen-CO-FISH experiment shown in G-H. (G) (Left) Representative Cen-CO-FISH on metaphase chromosomes after CENP-A depletion in the previous S-phase. Scale bar: 5 μm. (Right) Schematic of the resulted Cen-CO-FISH staining patterns in normal and abnormal centromeres, with visible SCE due to recombination and cross over events. (H) Quantification of the percentage of aberrant Cen-CO-FISH patterns/cell after CENP-A depletion in the previous S-phase. Histogram is the average of three independent experiments (depicted with different shapes) with n = 15 cells per condition. Error bars: standard error of the mean. Mann-Whitney test was performed on pooled single cell data of three independent experiments: ****p<0.0001. (I) Quantification of the percentage of aberrant Cen-CO-FISH patterns/cell with or without CENP-A depletion in the different phases of the cell cycle as indicated. For synchronization strategy see figure S1D. n = 15 cells per condition. Red lines represent the median. Mann-Whitney test: ****p<0.0001.

To establish the critical time when CENP-A disruption causes alpha-satellite instability, we induced CENP-A depletion at different stages of the cell cycle (Figure S1D, E) and monitored recombination at centromeres in the first mitosis using centromeric chromosome-orientation fluorescence *in situ* hybridization (Cen-CO-FISH; Fig. 1E) (32). CENP-A depletion upon G1 release, 8 h after G1 release (corresponding to S-phase entry), or thymidine release into S-phase, but not in G2 or metaphase-arrested cells, triggered severe centromere rearrangements (Figure 1F-I and S1F). These data demonstrate that absence of CENP-A in S-phase increases recombination at centromeres.

Since DNA replication stress can induce recombination, we assessed if replication fork dynamics were altered by CENP-A depletion using DNA combing for single molecule analysis (33). We performed sequential labeling of nascent DNA with nucleotide analogs, CldU and IdU (Figure 2A) during early S when mostly euchromatin is replicated (34), and in late S when centromeres are replicated (18, 19) as confirmed by BrdU-Immunoprecipitation (Figure S2A). A change in CIdU + IdU tracks length was used as indicator of a change in replication fork progression. Removal of CENP-A did not alter replication fork speed at 3 h after thymidine-release (Figure 2B-C). However, at 7 h, when fork speed accelerated in untreated cells as previously observed (35), fork speed was reduced upon CENP-A depletion (Figure 2B, C and S2B,C). These phenotypes were not due to perturbations in cell cycle progression, as CENP-A-depleted cells entered and progressed through S-phase with the same kinetics as untreated cells (Fig S2D, E).

**Figure 2.**
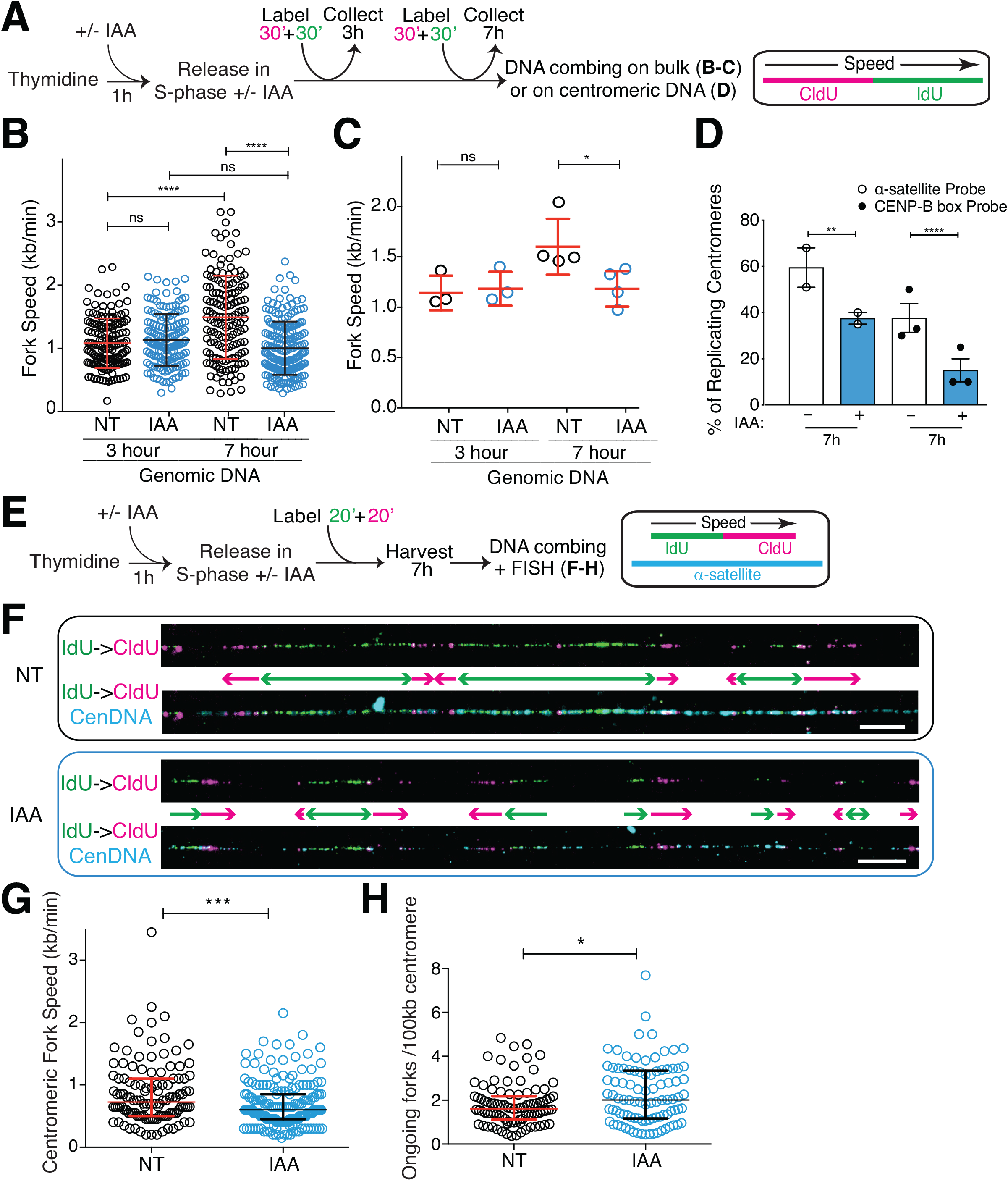
CENP-A removal leads to impaired replication fork progression through centromere alpha-satellites in late S-phase. (A) Schematic illustration of the single molecule DNA replication experiments shown in B-D. (B) Quantification of the replication fork speed in n>100 tracks/condition. The bars represent the standard deviation of the mean. Mann-Whitney test: ****p<,0.0001. (C) Summary of the quantification of the replication fork speed in the bulk genome at 3 h or 7 h thymidine release in 3 or 4 independent experimental replicates. The bars represent the standard deviation of the mean. Mann-Whitney test: *p=0.0286. (D) Quantification of the percentage of centromeres actively replicating DNA in the DNA combing assay (IdU and/or CldU-positive). Two different oligo PNA probes were used to target alpha-satellites. Each dot represents one experiment with n>30 centromeric tracks per condition. The bars represent the standard error of the mean. χ^2^ test: **p=0.005, ****p<0.0001. (E) Schematic illustration of the single molecule DNA replication experiments at centromeric region shown in E-G, where 6 RNA probes were used to target alpha-satellites. (F) Representative images of single molecule alpha-satellite DNA combing at 7 h thymidine release. A probe mix recognizing alpha-satellite repeats was used to specifically label centromeres. Scale bar: 10 μm. (G) Quantification of the centromeric replication fork speed at 7 h thymidine release. n>110 tracks/condition. The lines represent the median with interquartile range. Mann-Whitney test: ***p=0.0006. (H) Quantification of the number of ongoing forks/100 kb at centromeres at 7h thymidine release. n>90 tracks/condition. The lines represent the median with interquartile range. Mann-Whitney test: *p=0.0411.

To directly assess replication fork dynamics on the centromeric repetitive DNA, alpha-satellites were labelled by two different FISH probes on combed DNA. Among these FISH-positive segments of fibers ranging 50 – 200 kb, the overall frequency of IdU and/or CIdU labelled fibers at 7 h from thymidine release was reduced upon CENP-A depletion (Figure 2D), suggesting that replication at alpha-satellite repeats was impaired. To cover longer and more diverse stretches of alpha-satellites, we used a mix of 6 different ~50 bp probes that hybridized to alpha-satellites DNA ranging 50 – 1300 kb (median of ~250 kb) (Figure S2F). This analysis revealed that replication of centromeric DNA occurs in clusters (Figure 2E, F) leading to many converging forks at 7 h after thymidine release. As termination promotes topological burdens (36), this could contribute to slower replication at centromeres compared to the bulk genome (37). In CENP-A-depleted cells, replication fork speed at fibers labelled with the probes mix was reduced compared to untreated conditions (median 0.72 kb/min to 0.60 kb/min) (Figure 2G and S2G). Within replication clusters seen in late S-phase at centromeres (Figure 2F), we observed an increase in the number of ongoing forks/100 kb (median 1.60 vs. 2.01) upon CENP-A depletion (Figure 2H), indicating dormant origin firing, possibly reflecting compensation for the diminished fork velocity (Figure 2G). However, dormant origin firing was not sufficient to fully rescue timely replication of entire segments of centromeres (Figure 2D, F). Altogether, our data suggest that CENP-A is needed for efficient replication fork progression through alpha-satellite repeats.

As a candidate for the impediments that induce replicative stress at centromeres upon CENP-A depletion, we monitored the occurrence of DNA-RNA hybrids (R-loops), known to obstruct DNA replication fork progression (Figure 3A) (38). Immunofluorescence-based detection of DNA-RNA hybrids using S9.6 monoclonal antibody (39) showed that mean signal intensities at centromeres increase upon CENP-A removal only in late S-phase (Figure 3B-C and Fig. S3A), the time when most centromeres are replicated (Figure S2A). We confirmed the enrichment of centromere-specific DNA-RNA hybrids in absence of CENP-A in late S-phase using DRIP, which captures DNA-RNA hybrids in their native chromosomal context followed by RT-qPCR (Figure 3D-E). The signal detected with the S9.6 antibody was abrogated by treatment with RNase H (Figure 3B, D, E) confirming its specificity to DNA-RNA hybrids. Taken together, these data show that CENP-A depletion causes impaired DNA replication progression and R-loop formation at centromeres.

**Figure 3.**
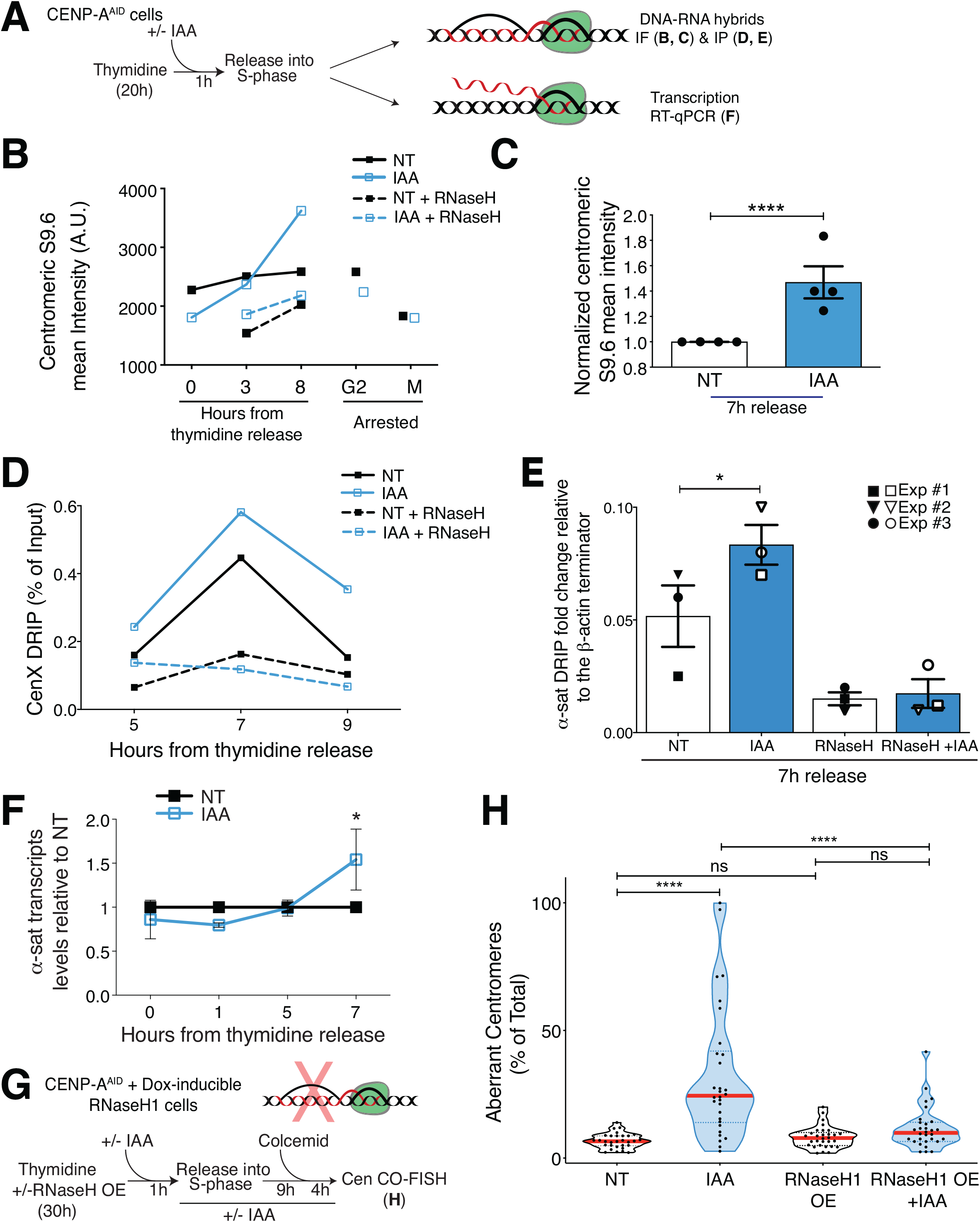
Accumulation of R-loops following CENP-A depletion causes increased centromere recombination. (A) Schematic illustration of the experiments shown in B-F. (B) Quantification of R-loops by mean fluorescence intensity of S9.6 antibody at centromeres upon CENP-A depletion through S-phase progression, in G2 or in mitotic cells. Lines represent a time course experiment. Mean of n>400 centromeres/condition. (C) Quantification of the R-loops mean fluorescence intensity at centromeres after 8h thymidine release. Each dot represents one independent biological replicate with n>300 centromeres per condition. The bars represent the standard error of the mean. A Mann-Whitney test was performed on the pooled single centromere data of four independent experiments: ****p<0.0001 (D) DRIP-qPCR quantification of the R-loops levels at the centromere of chromosome X at different timepoints in S-phase. n = 1. (E) DRIP-qPCR of R-loops levels at the centromeres using alpha-satellite primers at 7 h post-thymidine release. Each dot represents an independent experiment, depicted with a different shape. Paired t test: *p=0.0488 (F) Quantification of centromere transcripts levels using alpha-satellite primers by RT-qPCR after thymidine release. Each dot represents the mean of three independent experiments. The bars represent the standard error of the mean. Wilcoxon Signed Rank Test: *p=0.0312 (G) Schematic illustration of the rescue experiment shown in H. (H) Quantification of the percentage of aberrant Cen-CO-FISH patterns/cell after RNase H1 overexpression, and subsequent CENP-A depletion during the prior S-phase. n = 15 cells per condition. The red lines represent the median. Mann-Whitney test: ****p<0.0001.

R-loops, which form co-transcriptionally, are prevalent at replication-transcription conflicts (38, 40). Centromeres are actively transcribed throughout the cell cycle (17, 41), raising the hypothesis that CENP-A depletion causes R-loop formation due to convergence of replication and transcription machineries. Upon release from thymidine-mediated arrest, CENP-A depletion did not affect the levels of centromere transcripts during early S-phase, but when cells progressed to late S-phase, the RNA levels increased (Figure 3F). This increase was reproducible using different centromeric probes and across cell lines like U2OS and DLD-1 (Figure S3B). We also confirmed that *de novo* centromere transcription is indeed active in late S-phase by immuno-precipitation followed by RT-qPCR of 5-fluorouridine (FU) (Figure S3C-E).

Consistent with the notion that R-loops derive from transcription-replication conflicts (42), levels of γH2AX at centromeres in late S-phase increased after CENP-A depletion (Figure S3F-H). Furthermore, while the large majority of active RNA polymerase II dissociates from mitotic chromosomes (43, 44), an increase in centromere-associated active polymerase after CENP-A depletion can be detected during mitosis (Figure S3I, J). This increase was concomitant with the accumulation of the DNA damage kinase ATR (Figure S3I, J), in line with the previously reported R-loop-driven ATR activation at mitotic centromeres (21).

To test if R-loops could cause centromere damage and alpha-satellite repeat instability, we stably integrated RNase H1 under a doxycycline-inducible promoter in the hTERT-RPE-1 ^AID^CENP-A cell line (Figure 3G) (21). Expression of RNase H1 was able to rescue phenotypes observed in CENP-A-depleted cells such as enhanced centromere-associated γH2AX (Figure S3H) and aberrant Cen-CO-FISH patterns, indicative of centromere recombination (Figure 3H). These data demonstrate that the centromere instability in absence of CENP-A is caused by the generation of centromeric R-loops during alpha-satellite DNA synthesis.

Replicative stress is a main driver of genomic instability and favors the emergence of cancer. It has been shown that persistent replication intermediates in mitosis are processed *via* the error-prone mitotic DNA synthesis (MiDAS), prominently observed at common fragile sites (CFS) (45). To test if the replication stress imposed by CENP-A depletion in S-phase results in MiDAS, we depleted CENP-A in asynchronous cell culture for 10 h and monitored incorporation of EdU during the subsequent first mitosis (Figure 4A), assuming that these mitotic cells had gone through S-phase without CENP-A. We noticed an increase in cells containing EdU foci following CENP-A depletion, to a similar level of the DNA replication slowdown induced with low dose of aphidicolin (45) (Figure 4B). These EdU foci were found enriched at centromeric regions with Proximity Ligation Assay (PLA), indicating that a subclass of centromeres continues to be replicated outside S-phase (Figure 4C).

**Figure 4.**
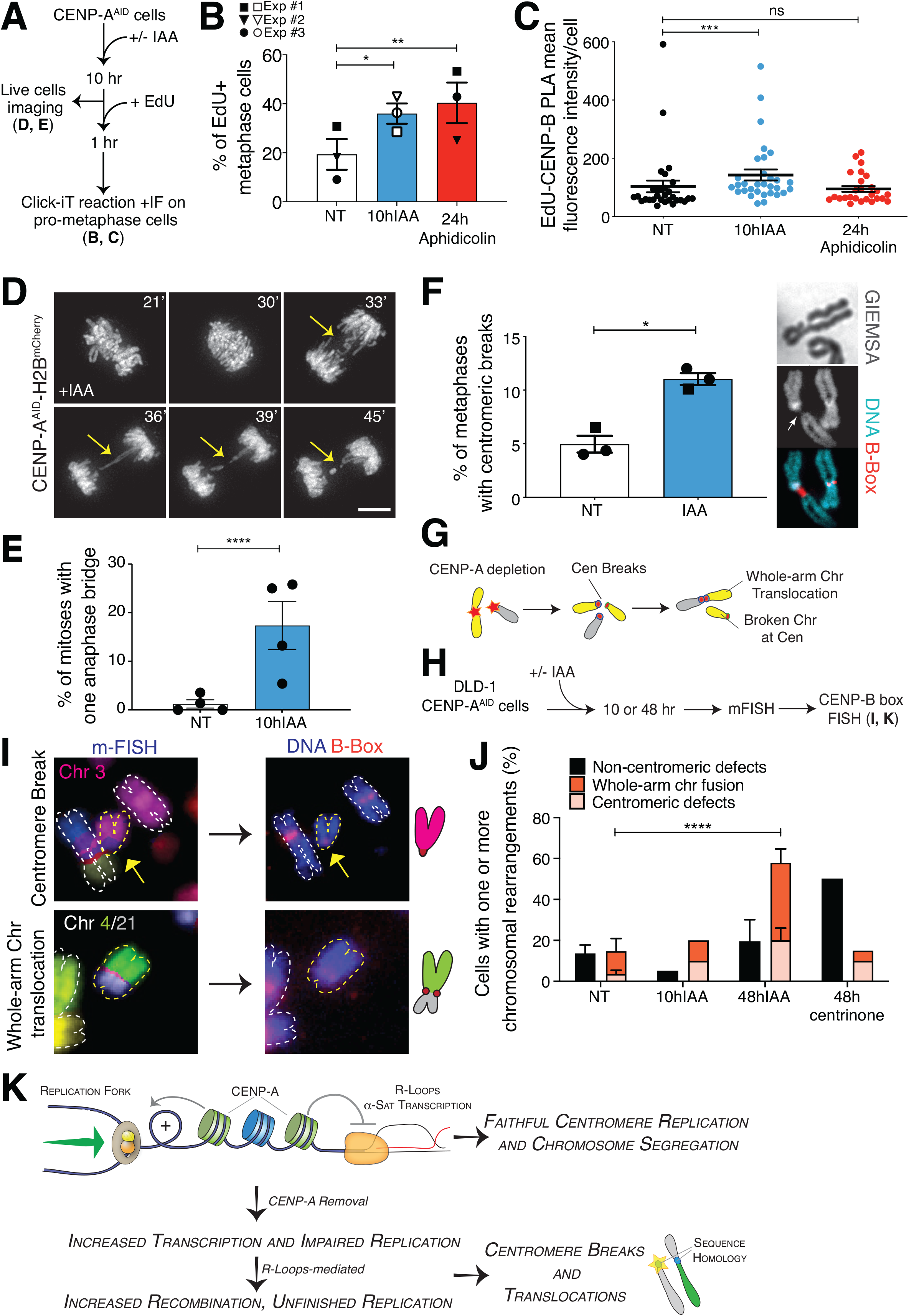
Centromere instability during DNA replication initiates chromosome rearrangements and translocations in the ensuing cell cycles. (A) Schematic illustration of the experiments shown in B-C. (B) Quantification of the percentage of EdU-positive metaphases after 10 h IAA or 24 h aphidicolin (0.2 μM). Each dot represents one experiment. The bars represent the standard error of the mean. χ^2^ test: *p=0.0435, **p=0.0044. (C) Quantification of EdU-positive centromeres in metaphase cells by measuring mean fluorescence intensity signal from PLA between CENP-B and EdU-biotin after 10 h IAA or 24 h aphidicolin (0.2 μM). n>25 cells/condition. The bars represent the standard error of the mean. Mann-Whitney test: ***p=0.0009. (D) Representative still images of H2B-mCherry RPE-1 live-cell imaging showing the presence of an anaphase bridge after 10 h IAA. Scale bar: 10 μm. (E) Quantification of the percentage of RPE-1 cells showing at least one anaphase bridge after 10 h IAA. Each dot represents one of four independent experiments. The bars represent the standard error of the mean. χ^2^ test: ****p<0.0001. (F) Quantification of the percentage of metaphase spreads showing reduced DAPI/Giemsa staining at the centromere after 24 h IAA (circle) or after thymidine release (square). Each dot represents one experiment with n>70 cells. The bars represent the standard error of the mean. χ^2^ test: *p=0.0113. (G) Illustration of the formation of centromeric chromosomal rearrangements after CENP-A depletion due to breaks generated during the previous cell cycle and repaired by non-allelic homologous recombination (NAHR). (H) Schematic representation of the multicolor-FISH experiments shown in I. (I) Representative images of mFISH and centromeric FISH (CENP-B box probe) of DLD-1 chromosome spreads showing chromosomal aberrations occurring at centromeres after 48 h IAA. A schematic of the observed translocation is also shown. (J) Quantification of the percentage of metaphase spreads showing chromosomal rearrangements by multicolor-FISH after CENP-A depletion or centrosome depletion in DLD-1 cells. N = 3 (NT & 48 h IAA) or 1 (10 h IAA & 48h centrinone 0.2 μM). The bars represent the standard error of the mean. χ^2^ test: ****p<0.0001. (K) Model for maintenance of centromeric DNA stability mediated by CENP-A-containing chromatin. See text for details.

Mitotic entry with under-replicated or unresolved recombination intermediates causes anaphase bridges (46). To monitor if replication stress induced by CENP-A depletion generates such mitotic perturbations, we used live cell imaging to follow chromosome separation in RPE-1 cells. CENP-A depletion in G1/early S (Figure 4A) indeed increased frequency of chromosome bridges threaded between the segregating DNA masses (Figure 4D, E and Suppl. Movie 1). In contrast, the number of ultrafine-bridges (UFBs) marked with PICH, often seen at centromeres (47), was not affected by CENP-A depletion (Figure S4A), implying different underlying molecular origins (48, 49) and substantiating that most centromeric UFBs are not originating from under-replication but from DNA decatenation impairment (50). Moreover, we detected a significant increase in regions with no or reduced DAPI/Giemsa staining – possibly due to chromosome breakage, under-replicated or decondensed regions – affecting one or more metaphase centromeres (Figure 4F) and resembling cytological abnormalities seen at CFS after replicative stress. We also found co-localization between the DNA double strand break factor 53BP1 and the centromeric sequence specific DNA binding protein CENP-B by PLA in the following cell cycle after CENP-A depletion (Figure S4B), suggesting that DNA breaks persist at the centromere in interphase. In summary, our data show that upon entry into mitosis, replication impairments caused by CENP-A depletion lead to centromere instability, therefore mimicking fragile sites and under-replicated regions of the genome.

DNA damage at centromere segments may induce chromosome translocations between chromosomes with homologous centromere sequences (Figure 4G). To score them, we combined multicolor FISH (mFISH) followed by CENP-B boxes FISH staining to check the status of centromeric DNA (Figure 4H). Within two cell cycles following CENP-A depletion, centromeric abnormalities and rearrangements were induced at one or more chromosomes, beyond numerical aneuploidy. These abnormalities spanned from centromere breakage and fragmentation, isochromosomes and a high proportion of whole-arm chromosome translocations involving both acrocentric or metacentric chromosomes (Figure 4I, J and S4C-E). No chromosomal translocations were detected in cells arrested in the first mitosis without CENP-A (10hIAA) (Figure 4J), indicating that anaphase bridge breakage is a necessary step for centromere instability to trigger overall genome instability, as recently shown for mitotic chromosome breakage-fusion-bridge (BFB) cycle (51). To ensure that this phenotype was not due to whole chromosome mis-segregation resulting from CENP-A depletion (27), we generated mitotic defects independently of centromeric dysfunction by centrosome depletion with a Plk4 inhibitor (48h centrinone) (52). In this case, we still detected an increase in chromosomal rearrangements, but such alterations occurred mainly outside the centromeric regions (e.g. telomere fusions; Figure 4J and Figure S4C-D) confirming the specific effect of CENP-A depletion on centromere DNA fragility and chromosome integrity. These results imply that chromosome translocations at centromeres are not the primary consequence of chromosome missegregation, but are generated by mitotic DNA breakages due to replication stress at centromeres.

## Discussion

Here we demonstrate that CENP-A is critical in the maintenance of centromeric DNA repeats by repressing R-loop formation during DNA replication (Figure 4K). While impaired histone assembly onto newly replicated DNA impedes replication (53), our CENP-A depletion did not interfere with histone assembly during S-phase (Figure S1B-C). Instead, CENP-A nucleosomes removal on parental DNA slows replication fork progression, likely due to replication-transcription conflicts, which stabilize R-loops. While nucleosomes generally suppress transcriptional initiation (54) and compete with R-loop formation (40), CENP-A depletion increased R-loop and centromeric RNA levels only during centromere replication. Thus, we propose that CENP-A chromatin holds a specialized function to facilitate fork progression and suppress R-loop formation at centromeres during DNA replication. Since alpha-satellite RNAs stay associated with the centromere from which they were transcribed throughout the cell cycle (17), CENP-A chromatin may sequester these RNAs to minimize R-loop formation upon replication-transcription conflicts. This is consistent with the report that CENP-C and other centromeric proteins can bind to RNA (17), and depletion of CCAN (Constitutive Centromere Associated Network) components (CENP-C, CENP-T, CENP-W) induces aberrant centromere recombination without obvious displacement of CENP-A from chromatin (10). It is also possible that CENP-A is important for recruitment of proteins, such as helicases that remove DNA-RNA hybrids (38). Altogether, our results suggest that centromeric DNA regions are intrinsically difficult to replicate to a similar extent as other repetitive sequences where specialized proteins act to facilitate fork progression (55, 56).

Mitosis with under-replicated DNA and/or recombination caused by CENP-A depletion would generate DNA damage to a subset of centromeres, which can promotes non-allelic exchanges between near-identical repeats on different chromosomes. Centromere clustering may facilitate chromosome translocation at centromeres (57). Notably, inter-centromeric rearrangements following centromere instability increased preferentially in acrocentric chromosomes, which are frequently linked together at their rDNA loci forming the nucleoli (58). As CENP-A levels are reduced in senescent cells (59) and in certain type of organismal aging in human (60), centromere-induced structural aneuploidy may represent a key mechanism underlying aging-associated tumorigenesis. Senescent cells present hypomethylated and highly transcribed centromeres (61, 62) that show altered nucleolar association (63) that in turn might impact the regulation of alpha-satellite expression (41) and favorize rearrangements at acrocentric chromosomes. During neocentromere formation, CENP-A can also be lost from the original alpha-satellite locus which has been reported to lead to centromere erosion at the repeats (64). Alpha-satellite fragilities may thus explain the high prevalence of centromeric transcripts overexpression and whole-arm chromosome translocations in cancer (11, 12).

## Methods

### Cell culture

Cells were cultivated at 37°C in a 5% CO2 atmosphere with 21% oxygen. Immortalized hTERT retinal pigmented epithelial (RPE-1) cells were grown in Dulbecco Modified Minimal Essential medium DMEM:F-12 (Life Technologies) supplemented with 10% fetal bovine serum (FBS), or 10% tetracycline-free FBS for RNase H1-GFP overexpression experiments (Atlanta Biologicals), 100 U/ml Penicillin-Streptomycin (P/S; Life Technologies), 0.123% sodium bicarbonate and 2 mM L-glutamine. Flp-In TRex DLD-1 cells and U2OS 2-6-3 R.I.K LacO cells were maintained in Dulbecco’s modified essential medium (DMEM) containing 10% FBS.

### Cell synchronization & treatments

In preparation to the experiment, hTERT RPE-1 cells were split at low confluency (~ 3 × 10^5^ cells/10 cm dish) approximately 24 h prior to the treatment. Indole-3-acetic acid (IAA, auxin) (I5148; Sigma) was dissolved in ddH2O and used at a final concentration of 500 μM. RNase H1-GFP expression was induced using 200 ng/μl of doxycycline. RNase H treatment on fixed cells was performed using 5 U RNase H (M0297, NEB) for 3 h at 37°C, or mock for no RNase H digestion control in 1X RNase H digestion buffer without the enzyme, before proceeding to immunofluorescence. Thymidine synchronization and release experiment was performed by incubating cells with 2-5 mM Thymidine for 22-28 h; cells were then treated with IAA for 1 h, washed three times with Dulbecco sterile phosphate-buffered saline (PBS) and released into fresh medium containing IAA. For G1 synchronization, cells were treated for 24 h with 150 nM Palbociclib (S1116; Selleck), a CDK4/6 inhibitor before washing out and releasing from the arrest – according to a protocol established and kindly shared by Eleanor W. Trotter and Iain Hagan. G2 arrest was induced by RO-3306 CDK1 inhibitor (ALX-270-463, Enzo) at 10 μM final concentration for 20 h. Mitotic arrest was achieved using 0.1 μg/mL Colcemid (Roche) for 5 h, followed by mitotic shake off to obtain a pure mitotic population for processing. Centrinone treatment to induce centrosome loss was performed at 0.2 μM for 48 h.

### Fluorescence in situ hybridization (FISH)

RPE-1 cells were released from a 24 h thymidine block into medium with or without auxin and trapped in mitosis with a 3 h treatment in colcemid. Cells were swollen in hypotonic solution (75 nM KCl) and fixed with acetic acid and spread onto slides. Coverslips were rinsed with 80% ethanol, air dried and a mix of CENP-B Boxes Cy3 (PNA, Bio) probe diluted in hybridization buffer (1:250) was applied. Probe and sample were denaturated by heating (72°C, 2 min) followed by overnight incubation at 37°C. Next day, slides were washed with 0.4X SSC at 72°C for 2 min, followed by a quick wash in 2X SCC, 0.05% Tween 20 at RT for 30 sec. Slides were rinsed with PBS, mounted in ProLong Gold antifade reagent with DAPI (P36935, Invitrogen) and imaged with an epifluorescence microscope (Leica DM6000).

### Cen-CO-FISH

CO-FISH at centromere was performed as previously described (*24*). Cells were grown in 10 μM BrdU:BrdC (3:1) for 16–20 h before incubation with 0.1 μg/mL colcemid (Roche) for 2-3 h. Harvested cells were incubated in pre-warmed 0.075 M KCl at 37°C for 30 min. Cells were then fixed in freshly made 3:1 methanol/acetic acid and dropped onto glass slides in a Thermotron Cycler (20°C, 50% humidity). Re-hydrated slides were treated with RNase A (0.5 μg/ml in PBS, Sigma R5000) for 10 min at 37°C, stained with 0.5 μg/ml Hoechst 33258 (Invitrogen) in 2X SSC for 10 min at RT and exposed to 365-nm UV light (Spectralinker XL-1000 1800 UV irradiator; Spectronics Corp.) for 30 min. The BrdU/C labelled DNA strand was digested with 10 U/ml Exonuclease III (Promega M1811) for two rounds of 10 min at RT, followed by dehydration washes with 75%, 95% and 100% ethanol. The slides were allowed to air dry fully before applying Hybridizing Solution (70% formamide, 1 mg/ml blocking reagent (Roche), 10 mM Tris-HCl pH 7.2) containing the PNA probe (CENP-B Box Cy3: ATTCGTTGGAAACGGGA; PNABio) and hybridized at RT for 2 h. The slides were washed briefly in wash buffer 1 (70% formamide/10 mM Tris-HCl) before the second 2 h hybridization with the complementary PNA probe (CENP-B Box RC Alexa488: TCCCGTTTCCAACGAAT; PNABio). Slides were then washed in buffer 1 for 30 min, followed by washes in 0.1 M Tris-HCl, pH 7.0/0.15 M NaCl/0.08% Tween-20. DNA was counterstained with 4,6-diamidino-2-phenylindole (DAPI; Sigma D-9542). Slides were mounted in antifade reagent (ProLong Gold, Invitrogen) and imaged.

### Chromatin extraction and immunoblotting

Cells were harvested by scraping into lysis buffer (250 mM NaCl, 50 mM Tris pH 8.0, 5 mM EDTA, 0.5% NP-40) supplemented with protease inhibitors (Roche, cat. # 1 697 498), phosphatase, Protein Phosphatase Inhibitor 2 (New England Biolabs) and Trichostatin A HDAC inhibitor (TSA; Sigma) into cold microcentrifuge tube and kept on ice for 15 min. Tubes were spun 10 min, 14000 RPM at 4°C to pellet chromatin. The soluble fraction in the supernatant was removed. Chromatin was resuspended in equal volume lysis buffer and sonicated at 4°C using Bioruptor ® (Denville, NJ) for 45 min. Concentration was measured using a NanoDrop 1000 spectrophotometer (Thermo Fisher Scientific) and samples were prepared into 2X Loading Buffer, and resolved by SDS-PAGE, transferred onto nitrocellulose membranes. Membranes were ponceau stained to ensure equal loading, blocked with Odyssey Blocking Buffer (Li-Cor 927-40000) and incubated with CENP-A Mouse antibody (1:500; Ab13939; Abcam) in 5% BSA TBS with 0.1% Tween 20 and detected with IRDye 800CW and 680LT secondary antibodies using the Odyssey Infrared Imaging System (Li-Cor).

### Histone deposition monitoring with SNAP-labelling

In order to label newly deposited histones, the old H4-SNAP pool was first quenched with 3 μM BTP (NEB) for 30 min in complete medium, then cells were washed two times with fresh medium, after which cells were reincubated in complete medium to allow excess compound to diffuse from cells for 30min and washed again two times with fresh medium.

H4-SNAP histones were labelled with 3 μM TMR fluorophore (NEB) for 30 min in complete medium, then washed two times with fresh medium. After 30 min of recovery, cells were washed again two times with fresh medium and then fixed for immunofluorescence.

### Generation of Stable Cell Lines

4 μg of lentiviral plasmid (pINDUCER20-RNaseH-GFP) was co-transfected into a 10 cm dish of 293-T cells with 4 μg psPAX2 and 4 μg pMD2.G plasmids using Fugene HD transfection reagent for the production of lentivirus. The resulting lentiviral supernatant was mixed with 8 μg/mL polybrene and incubated with RPE-1 cells for 24 hr. Stable integration was selected with 10 μg/ml blasticidin S for H4-SNAP and 0.4 mg/mL G418 for RNaseH1-GFP. H4-SNAP single clones were isolated after serial dilutions.

### Flow cytometry

For cell cycle and BrdU analyses, cells were processed using the Cell Cycle and BrdU Flow Kit (BD Biosciences) according to manufacturer’s instructions. Briefly, the cells were fixed, permeabilized and stained using BrdU antibody directly conjugated with a FITC secondary. DNA was stained using DAPI. Data were acquired using a BD LSR I Flow Cytometer, and processes using FlowJo software v9.

### EdU staining in metaphase cells

Metaphase cells were enriched by 2 h treatment with colcemid, EdU (10 μM final) incorporation was performed in the last hour before fixation. EdU click labeling was performed using Click-iT® labelling technologies (Thermo-Fisher scientific) or a homemade click chemistry buffer (Tris-HCl 100 mM, pH 8.5, CuSO4 1mM, Azide-fluor 488/Biotin azide 5 μM, Ascorbic acid 100 mM) for 30 min at room temperature on fixed and permeabilized cells.

### Immunofluorescence, Proximity Ligation Assay (PLA) and Antibodies

Cells were grown on poly-L-lysine-coated coverslips, and fixed in 2-4% formaldehyde at room temperature for 10 min. Pre-extraction with 0.1% Triton-X in PBS for 45sec prior to fixation or permeabilization with 0.2% Triton X-100 in PBS for 5 min after fixation were applied depending on the antibody. After blocking in 5% Bovine Serum Albumin in 0.1% Tween-20 in PBS for 10 min or in 2.5% FBS (v/v), 0.2 M Glycine, 0,1% triton X-100 (v/v) in PBS for 30 min at room temperature. Primary antibodies were incubated in blocking buffer for 1 h at room temperature. Cells were washed three times with PBS and incubated with secondary fluorescent antibodies (1:500/1:1000) for 45 min at RT. The following antibodies were used: 53BP1 (1:500; NB100-304, Novus Biologicals), ACA (1:500; 15-235-0001, Antibodies Incorporated), CENP-A mouse (1:500; ADI-KAM-CC006-E, ENZO), CENP-A rabbit (1:200; #2186; Cell Signaling), RNA-DNA Hybrid clone S9.6 (1:200; MABE1095; Millipore), CENP-C (1:1000; PD030, Clinisciences), Biotin (1:2000; ab53494, Abcam), ACA (1:500; 15-235-0001, Antibodies Incorporated), γH2AX (1:200; 05-636 Millipore), SMARCAL1 (1:200; Ab154226; Abcam). All secondary antibodies used for immunofluorescence microscopy were purchased from Jackson Immuno Research. After immunostaining, cells were DAPI counterstained and mounted using anti-fading reagent Prolong Gold (Life Technologies). Proximity Ligation Assay was performed using the Duolink® In Situ PLA reagents from ThermoFisher following the manufacturer’s instructions.

### Ultrafine bridges IF staining

Asynchronous RPE-1 cells grown on coverslips were fixed using Triton X-100-PFA buffer (250 mM HEPES, 1X PBS, pH7.4, 0.1% Triton X-100, 4% methanol-free paraformaldehyde) at 4 °C for 20 min, then washed 5 times with 1x PBS. Cells were permeabilized with permeabilization buffer (0.5% Triton X-100, 1× PBS) for 20 min on ice, then washed 5 times with 1X PBS. Cells were blocked with serum (0.5%FBS in 1×PBS) at room temperature for 15min followed by primary antibodies incubation at 37°C for 1.5hr in a humid chamber. Next, cells were washed 5 times with 1X PBS and secondary antibody incubation was performed for 30min at room temperature followed by 5 washes with 1× PBS and coverslips mounting.

The following antibodies were used: ACA (1:500; 15-235-0001, Antibodies Incorporated) and PICH (1:200; H00054821-B01P, Abnova).

### Microscopy, Live-Cell Microscopy and Image Analyses

Images were acquired on a Fluorescent microscope DeltaVision Core system (Applied Precision) with 100X Olympus UPlanSApo 100 oil-immersion objective (NA 1.4), 250W Xenon light source equipped with a Photometrics CoolSNAP_HQ2 Camera. ~ 4μm Z-stacks were acquired (Z step size: 0.2 μm). Imaris software (Bitplane) was used to quantify fluorescence intensity in the deconvolved 3D images using centromere surfaces automatically determined by ACA staining detection. Quantification of structured illumination (SI) images at centromeres was performed using Imaris software (Bitplane) surface fitting function and extracting data on each centromere volume and sphericity. All images presented were imported and processed in Photoshop (Adobe Systems, Inc S.J.). Movies of live cells were acquired using an Inverted Eclipse Ti-2 (Nikon) full motorized + Spinning disk CSU-W1 (Yokogawa) microscope. Cells were grown on high optical quality plastic slides (ibidi) for this purpose.

### Metaphase spreads centromere quantification

In order to measure the fluorescence intensity of proteins at metaphase-arrested centromeres, points were manually chosen outside the two centromeres of each individualized chromosome. A line was then drawn between the two points and the fluorescence along the line was measured for each channel using the Plot profile function in Fiji. Background correction was performed by subtracting the lowest pixel intensity value to the intensity values along the line. The line length was normalized in order to compare and pool data coming from different chromosomes; the fluorescence intensity values were extrapolated accordingly.

### Unfixed chromosome spreads

RPE-1 cells were grown to 80% confluency on coverslips and treated with colcemid for 2 h. Cells were incubated with hypotonic medium (60% medium, 40% water) for 3 min at 37°C and then centrifuged at 2500 rpm for 10 min. Cells were blocked in KCM buffer (120mM KCl, 20mM NaCl, 10mM Tris-HCl pH=7.7, 0.1% Triton-X-100, 0.5 mM EDTA) + 1% BSA for 30 min. Incubations with primary antibodies were conducted in blocking buffer for 1 h at room temperature using the following antibodies: ACA (1:500; 15-235-0001, Antibodies Incorporated), ATR (1:1000; A300-138A, Bethyl Laboratories) and RNA polymerase II S2P (1:1000; ab24758, Abcam). Samples were washed in KCM three times and then incubated with the respective secondary antibody (1:500) in blocking buffer for 45 min. Cells were washed in KCM three times and then fixed in KCM+4% formaldehyde for 10 min prior to DAPI staining and slide mounting. Images were acquired on a Fluorescent microscope DeltaVision Core system (Applied Precision) with 100X Olympus UPlanSApo 100 oil-immersion objective (NA 1.4), 250W Xenon light source equipped with a Photometrics CoolSNAP_HQ2 Camera. 4 μm Z-stacks were acquired (Z step size: 0.2 μm).

### DNA combing

Plug preparation: RPE-1-AID-CENP-A cells with/without auxin were pulse labeled with CldU (25 μM) for 30 min, washed 2 times with warm media, followed by pulse labeling with IdU (50 μM) for 30 min. Cells (2.5 × 10^5^) were harvested and resuspended in 1X PBS-NaN3 (0.2%) and embedded in 2% low-melt agarose (Mb grade, BioRad). Agarose plugs were chilled at 4°C for 30 min and then ejected into Proteinase K digestion buffer (1 mg/mL proteinase K, 1% N-Lauroylsarcosine, 0.2% sodium deoxycholate, 100 mM EDTA, 10 mM Tris-HCl, pH 8.0) and incubated overnight at 55°C. Plugs were then washed at least thrice in 1X TE buffer (pH8.0) containing 100mM NaCl for at least 1 h. Plugs were then melted in 50mM MES buffer (pH 5.5) at 68°C for 20 min and transferred to 42°C where 1.5 μL of ß-agarase (New England Biolabs) was added and incubated overnight. DNA solution was then poured into Teflon reservoirs and combed onto silanized coverslips using the Molecular Combing System (Genomic Vision) at a fixed speed of 300 μm/s to give a stretching factor of 2 kb/μm. Coverslips were made in house using a previously established method (65). Note that IdU and CldU were incorporated for 20 min each to measure replication at the centromere.

Immunofluorescence (bulk genome): For IF staining, slides were dried at 65°C for 2 h and then denatured in 0.5N NaOH +1M NaCl for 8 min, then dehydrated in ethanol series for 5 min each (70%, 90%, 100%). Slides were incubated in blocking buffer (5% BSA, 0.5% Triton-X100, 1X PBS) for 1 h at room temperature. Slides were incubated with anti-CldU (1:40, AbCam #6326) and anti-IdU (1:10, BD Biosciences #347580) in blocking buffer for 1 h at 37°C. Slides were washed 3X with PBS-Tween 20 0.5% and then incubated with secondary antibodies anti-mouse AlexaFluor 488 and anti-rat AlexaFluor 594. Slides were washed 3 times, and then blocked, and primary anti-ssDNA was incubated in block solution for 30 min at 37C. Slides were washed 3X, then incubated with anti-mouse AlexaFluor 647.

Immunofluorescence +FISH (centromere): The alpha-satellite probe mix consists of 6 oligos of ~ 50 bp each for a total of 321 bp. Oligos were designed to match the consensus sequence of 21 Higher Order Repeats families, distributed across all human centromeres, which exhibit CENP-A binding properties (8, 66).

CAGAAACUUCUUUGUGAUGUGUGCAUUCAACUCACAGAGUUGAACCUUUCUUUUGA;ACCUUUCUUUUGAUAGAGCAGUUUUGAAACACUCUUUUUGUAGAAUCUGCAA GUGG;AAGUGGAAAUCUGCAAGUGGAUAUUUGGACCGCUUUGAGGCUUUCGUUGGAAACGG;UAGACAGAAGCAUUCUCAGAAACUUCUUUGUGAUGUUUGCAUUCAACUCACAGAGU;GAACAUUCCCUUUGAUAGAGCAGGUUUGAAACACUCUUUUUGUAGUAUCUGCAAGU;GACAUUUGGAGCGCUUUGAGGCCUAUGGUGGAAAAGGAAAUAUCUUCCCAUAAAAA. DNA was denatured for 5 min in 1 N NaOH, followed by PBS (4°C) wash and dehydration in increasing concentrations of ethanol (75, 80 and 100%). Slides were hybridized overnight at 37°C with a biotinylated RNA alpha-satellite probe and the next day washed (3 times) with 50% formamide solution at RT. After 3 washes in 2X SSC and a quick wash in PBS, slides were incubated for 1h in blocking solution (blocking reagent Roche, 11096176001) at 37°C. Centromere signal was detected by alternating layers of avidin FITC (1:100, 434411, Thermo) and rabbit anti-avidin biotin conjugated (1:50, 200-4697-0100, Rockland laboratories) antibodies and replication was detected with rat anti-BrdU (1:25, ab6326, Abcam) and mouse anti-BrdU (1:5, 347580, BD Bioscience) for 1 h at 37°C. After a short wash step (0.5 M NaCl, 20 mM Tris, pH 7.8, and 0.5% Tween 20), the fibers were incubated with goat anti-mouse Cy5.5 (1:100, ab6947, Abcam) and donkey anti-rat 594 (1:50, 712-585-150, Jackson laboratory) for 20min. Fibers were mounted in ProLong Gold antifade reagent (P36935, Invitrogen) and acquisition was performed with an epifluorescence microscope (Apotome, Zeiss) with available mosaic option.

Alternatively, after dehydration combed slides were incubated with PNA-Biotin probes (alpha-satellite: TGATAGCGCAGCTTTGACAC; CENP-B box: ATTCGTTGGAAACGGGA) at 1:100 from 100μM stock in formamide at 42°C overnight. Slides were then washed twice for 15 min in 2X SSC, then incubated in blocking solution for 30 min. Slides were then incubated with Streptavidin Alexa 488 (1:50) for 20 min followed by a wash with PBS-Tween20 0.5% and 20 min incubation with anti-avidin followed by a wash with PBS-Tween20 0.5%. Streptavidin and anti-avidin sandwiches were repeated a total of 7 times. Thymidine analog staining was completed as above with secondary antibodies being anti-mouse Alexa Fluor 647 (IdU) and anti-mouse AlexaFluor 405 (ssDNA). Slides were mounted and imaged using an Olympus IX-71 DeltaVision Image Restoration Microscope (Applied Precision) and replication tracks measured using FIJI software.

### mFISH karyotyping

Cells grown to ~ 75–80% confluency were treated with colcemid for 3 h and prepared as previously described (8) for the mFISH karyotyping. The Metafer imaging platform (MetaSystems) and the Isis software were used for automated acquisition of the chromosome spread and mFISH image analysis.

### CUT&RUN-qPCR

CUT&RUN was performed according to the procedure previously reported (67) starting from 1 million cells and using anti-H4K5Ac (Abcam, ab51997, 1/1000) antibody. Rabbit IgG isotype control antibody (ThermoFisher, 10500C, 1/100) was used for background detection. qPCR was performed using the LightCycler 480 (Roche) system with primers reported below. Fold enrichement at centromeres was calculated using alpha-satellite primers, with the ΔΔCt method. The rabbit IgG isotype sample was used as control sample and normalization was performed to the Ct values from the Alu repeat primers.

### RT-qPCR

RNA was extracted from 1 million cells with RNeasy mini Kit (Qiagen) following the manufacturer’s instructions and with DNase treatment on column. In order to get rid of the remaining centromeric DNA contaminants, RNA was further digested with TURBO DNase overnight at 37°C (10 U per 2 μg of RNA) and purified with the RNeasy MinElute Cleanup Kit (Qiagen) following the manufacturer’s instructions.

Reverse transcription was performed on 2μg RNA with the High-Capacity cDNA Reverse Transcription Kit (ThermoFisher) following the manufacturer’s instructions. Each sample was also incubated without the reverse transcriptase enzyme in order to control for centromeric DNA contamination. The recovered cDNA was quantified by qPCR using a LightCycler 480 (Roche).

### qPCR primers

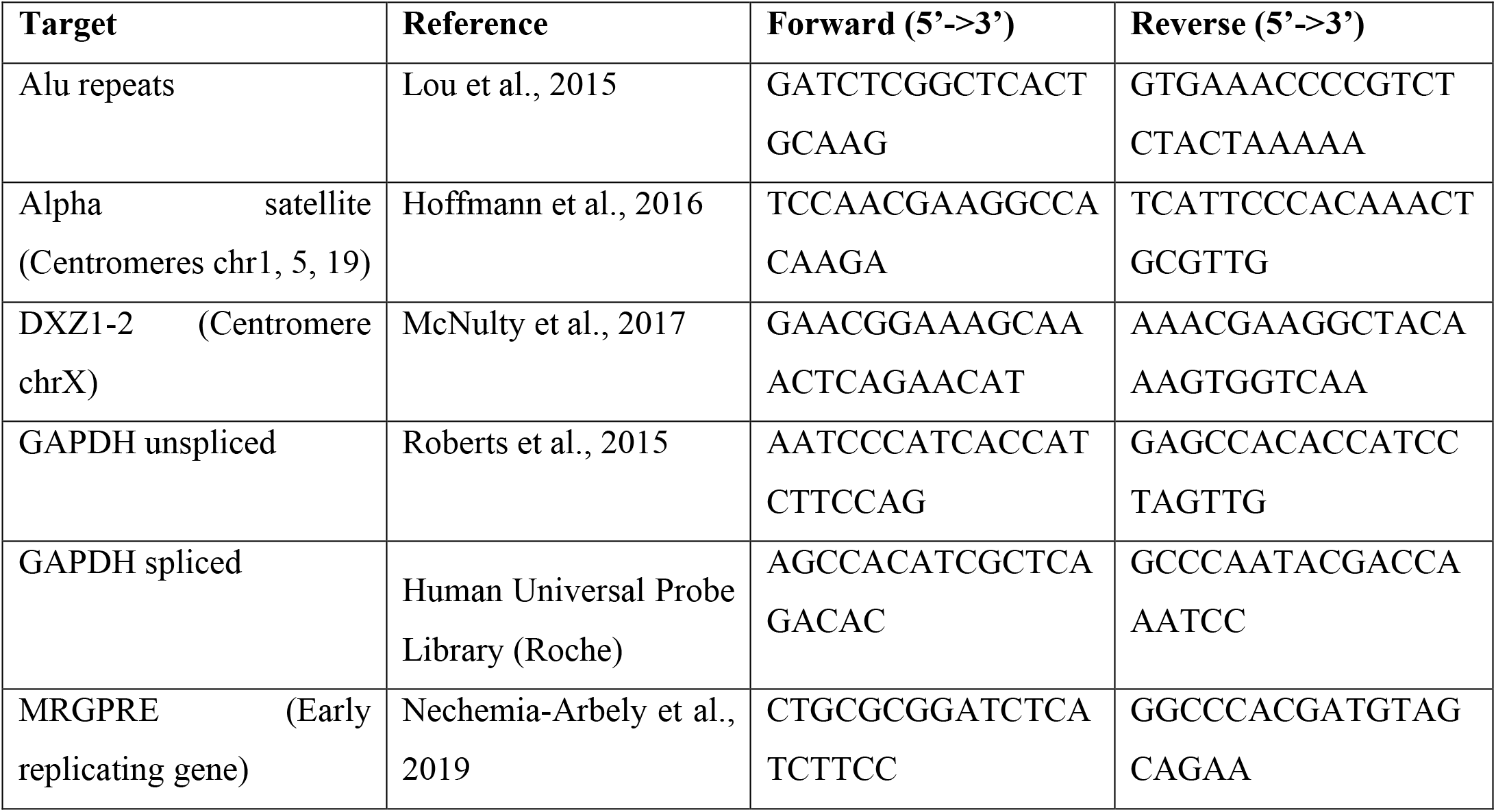

### 5-Fluouridine Immunoprecipitation for nascent RNA

This protocol is based on a previously described protocol of nascent transcripts capture (69). RNA was extracted from 5 million cells with RNeasy mini Kit (Qiagen) following the manufacturer’s instructions and with DNase treatment on column. In order to get rid of the remaining centromeric DNA contaminants, RNA was further digested with TURBO DNase overnight at 37°C (10 U per 2 μg of RNA) and purified with the RNeasy MinElute Cleanup Kit (Qiagen) following the manufacturer’s instructions.

25 μL protein G Dynabeads were coated with 2 μg of anti-BrdU antibody (#555627, BD Biosciences) for 10 min at room temperature and then blocked for 30 min at room temperature in blocking buffer (0.1% PVP, 0.1% UltraPure BSA, 0.1%Tween-20 in PBS). After two washes (0.1%Tween-20 & RNasin in PBS), beads were incubated with 2-4μg of pre-denaturated RNA (65 °C for 5 min) overnight at 4°C. Beads were washed three times and reverse transcription was directly performed on the beads with the High-Capacity cDNA Reverse Transcription Kit (ThermoFisher) following the manufacturer’s instructions. The recovered cDNA and the input cDNA were quantified by qPCR using a LightCycler 480 (Roche).

### BrdU-IP for nascent DNA & DNA-RNA hybrids Immunoprecipitation (DRIP)

DNA was extracted from 5 million cells with NucleoSpin Tissue Kit (Macherey Nagel) following the manufacturer’s instructions. The DNA was eluted in 130 μl TE and fragmented by sonication using a Covaris S220 sonicator to get 300-800 bp fragments with the following program:

Duty Factor: 5%

Peak Incident Power (W): 105 Cycles per Burst: 200

Time (seconds): 80

After setting aside 1% for input control, 2 μg of DNA were used for immunoprecipitation with 400 mL Binding buffer (10 mM NaPO4, pH 7.0/140 mM NaCl/0.05% Triton X-100) and 5 μg of the S9.6 antibody (Kerafast, 1mg/ml) or 5 μg of the BrdU antibody (#555627, BD Biosciences) on a rotative shaker at 4°C overnight.

At the end of the incubation, 25 μl of protein A magnetic beads prewashed 2−3 times with Binding buffer were added to the DNA/antibody complex and incubated for at least 4h at 4 °C on a rotative shaker. After 4 washes with Binding buffer, the beads were eluted with 250 μl elution buffer (50 mM Tris pH 8.0, 10 mM EDTA, 0.5% SDS) containing 7 μl proteinase K (20 mg/ml) for 45 min in a thermomixer at 55 °C at 800 rpm. Finally, the eluted DNA was purified using the NucleoSpin Gel and PCR Clean-up Kit (Macherey Nagel) and eluted in 50 μl TE for analysis by qPCR.

## Acknowledgments

We thank Mathieu Maurin (I. Curie) for helping with image analysis, Titia de Lange, Kaori Takai, Francisca Lottersberger, Nazario Bosco and other members of the de Lange Laboratory (Rockefeller U.) for helpful assistance with the CO-FISH technique, Iain Hagan and Wendy Trotter (U. of Manchester) for sharing synchronization information for Palbociclib, Florence Larminat (IPBS) and Kok-Lung Chan (Sussex U) for suggestions on DRIP and UFBs respectively, Hai Dong Nguyen & Lee Zou (Massachusetts General Hospital Cancer Center), Brooke Conti (Rockefeller U.), Lars Jansen (Oxford U), Renata Basto (I. Curie) for sharing reagents, and Jennifer Gerton, Karen Miga, Tatsuo Fukagawa and Tetsuya Hori for sharing unpublished data. We also thank the Flow cytometry platform, the Cell and Tissue Imaging facility (PICT-IBiSA, member of the French National Research Infrastructure France-BioImaging ANR10-INBS-04), the antibody facility platform and the sequencing platform at Institut Curie, Bio-Imaging Resource Center and Flow Cytometry Resource Center at the Rockefeller University.

## Authors contribution

D.F., H.F., S.G. and S.H. designed the study, made figures and wrote the manuscript. A. Scelfo. and R.G. designed the centromere probe mix. R.W., A. Smogorzewska, T.W. and S.G. conducted the single molecule replication assays. M.D., G.R., C.K.W. and D.F. performed the mFISH experiments. S.G. and S.H. performed and analyzed all other experiments.

