## Supplemental Text and Figures for "CENP-A chromatin prevents replication stress at centromeres to avoid structural aneuploidy"

**Contains:**

Supplementary Figure Legends S1-S4

Supplementary Figures S1-S4

Movies S1

#### Supplementary Figure Legends

**Figure S1. Rapid and timed chromatin disruption at human centromeres.** (A) Chromatin immunoblot showing $^{EYFP-AID}$CENP-A protein levels in asynchronous (A/S) or thymidine-synchronized cells following treatment with auxin (IAA) for 1 h. Ponceau staining was used as a loading control. The asterisk marks a non-specific band. (B) Quantification of the TMR mean centromeric fluorescence intensity used to label the new histone H4 pool after CENP-A depletion from G1 into early S cells (5 h IAA, quantification done on early S EdU+ cells) or during S-phase (7 h IAA, quantification done on late S EdU+ cells). Each dot represents one experiment with n>30 cells per condition. The bars represent the standard error of the mean. A Mann-Whitney test was performed on the pooled single centromere data of the three independent experiments: ****p<0.0001. (C) Quantification of the newly deposited nucleosomes along the cell cycle by H4K5Ac CUT&RUN-qPCR at the centromeres using α-satellite primers. Data were normalized to an IgG control sample and to Alu repeats. The bars represent the standard error of the mean. Lines represent a time course experiment. A schematic of the experimental setting is also shown. (D) Schematic illustration of the cell synchronization and CENP-A depletion strategy for Cen-CO-FISH experiments in Fig.1I. For CENP-A depletion in G1, cells were synchronized by CDK4/6 inhibitor palbociclib (24 h, 150 nM) and released in the presence of IAA for 10 h, then colcemid was added for 4 h to enrich for metaphase cells. For CENP-A depletion in S-phase, cells were synchronized palbociclib (24h, 150 nM) and released into fresh medium. IAA was added 8 h after release (corresponding to mid-S) then kept for an addition 10 h with colcemid added for 4 h to enrich for metaphase cells. For CENP-A depletion in G2, cells were synchronized at the G2/M transition by CDK1 inhibitor RO-3306 (24 h, 10 μM) and IAA was added 1 h before release into colcemid-containing medium for 4 h to enrich for metaphase cells. For CENP-A depletion in mitosis, cells were synchronized in metaphase by colcemid (5 h, 0.1 mg/ml) followed by mitotic shake-off. Cells were replated in IAA and colcemid-containing medium for 5 h. (E) Representative FACS plots showing the efficiency of cell synchronization in RPE-1 cells. (F) Single experiment data (same as Fig.1H) of the quantification of the percentage of aberrant Cen-CO-FISH patterns/cell after induction of CENP-A depletion in the previous S-phase with n=15 cells per condition. The lines represent the median. Mann-Whitney test: Exp1 **p=0.0024, Exp2 *p=0.0478, Exp3 **p=0.0012.

**Figure S2. Fork dynamics at late-replicating alpha-satellite DNA are affected by CENP-A depletion.** (A) Quantification of BrdU enrichment at the centromeres by BrdU-Immunoprecipitation-qPCR using alpha-satellite primers to assess replication timing in RPE-1 cells after thymidine release. The early replicating gene MRGPRE was used as control. n=3 or 2. The bars represent the standard error of the mean. (B) Representative images of bulk genome DNA combing at 3 h and 7 h thymidine. Scale bar: 20µm. (C) Quantification of the first replication track speed (CldU) in fibers positive for both CldU and IdU. This quantification eliminates the influence of the high number of termination events at centromeric replication clusters in the estimation of the fork speed. n>100 tracks/condition. The bars represent the standard deviation of the mean. Mann-Whitney test: ****p<0.0001. (D) Representative FACS plots showing the efficiency of RPE-1 cell synchronization by thymidine. (E) Representative FACS plots showing the amount of RPE-1 cells actively replicating DNA with BrdU incorporation at 7 h thymidine release. (F) Quantification of the centromeric signal length in kilobases (Kb) in the indicated condition. (G) Quantification of the centromeric first replication track speed (IdU) at 7 h thymidine release in fibers positive for both IdU and CldU. This quantification eliminates the influence of the high number of termination events at centromeric replication clusters in the estimation of the fork speed. n>110 tracks/condition. The bars represent the median with interquartile range. Mann-Whitney test: **p=0.035.

**Figure S3. CENP-A depletion causes increased centromeric transcription and R-loops formation during centromere replication.** (A) Representative immunofluorescence images of DNA-RNA hybrids at centromeres at 7 h thymidine release in RPE-1 $^{EYFP\text{-}AID}$CENP-A using the S9.6 antibody. Scale bar: 10 µm. (B) Quantification of the transcript levels at the centromeres using alpha-satellite and Cen X primers and by RT-qPCR after 7 h thymidine release in DLD-1 and U2OS cells. Each dot represents the mean of one experiment. The bars represent the standard error of the mean. U2OS cells: Wilcoxon Signed Rank Test: *p=0.0312. DLD-1 cells: One sample t test: *p=0.0273 (C) Schematic representation of the experiments shown in D-E. (D) Quantification of the nascent GAPDH transcripts by 5-Fluorouridine-IP-qPCR at 7 h thymidine release in RPE-1 cells. The unspliced (immature) transcripts are enriched after IP. n=6. The bars represent the standard error of the mean. Mann-Whitney test: **p=0.0022. (E) Quantification of the nascent centromeric transcripts by 5-Fluorouridine-IP-qPCR at 7 h thymidine release in RPE-1 cells. n=3. The bars represent the standard error of the mean. (F) Schematic representation of the experiments shown in G-H. (G) Representative immunofluorescence images of DNA damage at centromeres at 7 h thymidine release in RPE-1 $^{EYFP\text{-}AID}$CENP-A using the double-strand break marker γH2AX. Scale bar: 10 µm. (H) Quantification of the centromeric γH2AX mean fluorescence intensity from thymidine release in the indicated conditions in RPE-1 cells. Each dot represents the mean of one experiment. Each replicate is depicted with a different shape. The bars represent the standard error of the mean. A Mann-Whitney test was performed on the pooled single centromere data of the three independent experiments: ****p<0.0001. (I) Representative immunofluorescence images of active RNA polymerase II (S2P) and ATR at centromeres on metaphase spreads. Scale bar: 2 µm (J) Quantification of the centromeric active RNA polymerase II (S2P) and ATR mean fluorescence intensity on metaphase spreads after 10 h IAA in RPE-1 cells. Each dot represents the mean of one experiment with n>100 centromeres/condition. The bars represent the standard error of the mean. A Mann-Whitney test was performed on the pooled single centromere data of several independent experiments: ****p<0.0001.

**Figure S4. Chromosome fragility and structural aneuploidy arise at centromeric regions following CENP-A removal during DNA replication.** (A) Representative immunofluorescence images of ultrafine anaphase bridges stained by PICH and centromeres (ACA). Scale bar: 5 µm. Quantification of the presence of ultrafine anaphase bridges in RPE-1 and U2OS cells after 12 h IAA. Each dot represents one experiment with n>30 cells per condition. The bars represent the standard error of the mean. Wilcoxon Signed Rank Test: p=0.125. (B) Representative immunofluorescence images of Proximity Ligation Assay (PLA) between CENP-B and 53BP1. Quantification of centromeric DNA damage in RPE-1 cells by measuring PLA mean fluorescence intensity signal between CENP-B and 53BP1 after 24 h IAA or 48 h IAA. n>65 cells/condition. The bars represent the standard error of the mean. Mann-Whitney test: ****p<0.0001. Scale bar: 5 µm. (C) Representative multicolor FISH (mFISH) karyotypes of pseudo-diploid DLD-1 cells in untreated condition or after 48 h of IAA or 48 h centrinone. The yellow arrows indicate centromeric chromosomal rearrangements. (D) Representative images of mFISH and centromeric FISH (CENP-B box probe) of DLD-1 chromosome spreads showing chromosomal aberrations occurring at centromeres after 48 h IAA (left) and non-centromeric aberrations after 48 h centrinone (right and within a red square). (E) Representative images of mFISH in DLD-1 cells showing whole-arm chromosome translocations after three rounds of 48 h IAA treatment.

**Movie S1. CENP-A depletion in S-phase leads to anaphase bridges in the subsequent mitosis.** Representative video of anaphase bridge formation in a RPE-1 cell expressing H2B-mCherry and treated with auxin for 10 h. Images were acquired every 3 min.

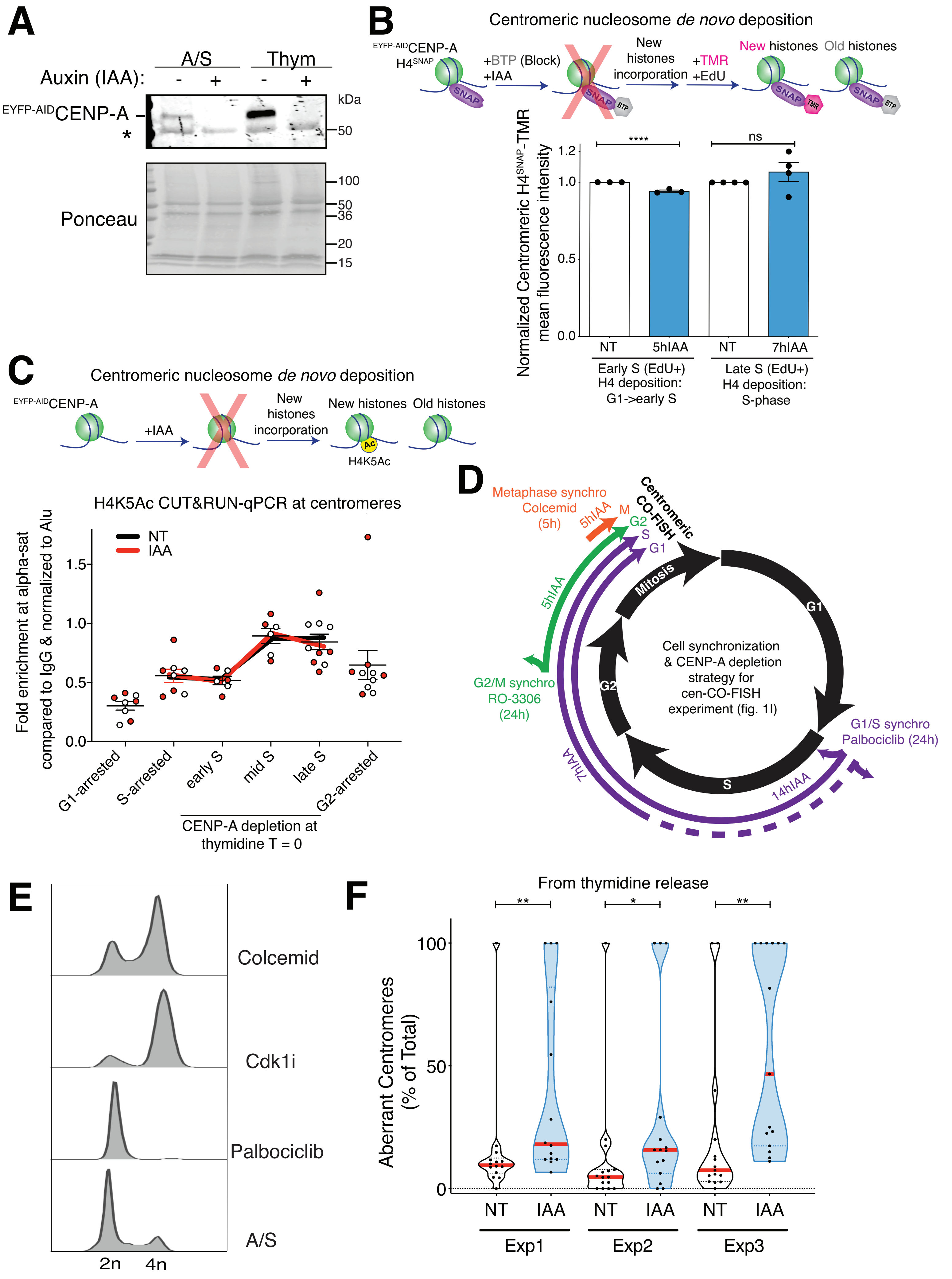
A
A/S
Thym
Auxin (IAA):
-
+
-
+
EYFP-AIDCENP-A
*
kDa
50
100
50
36
20
15
Ponceau
B
Centromeric nucleosome de novo deposition
EYFP-AIDCENP-A
H4SNAP
+BTP (Block)
+IAA
New histones incorporation
+TMR
+EdU
New histones
Old histones
SNAP
BTP
TMR
Normalized Centromeric H4SNAP-TMR mean fluorescence intensity
1.2
1.0
0.5
0.0
****
ns
NT
5hIAA
NT
7hIAA
Early S (EdU+) H4 deposition: G1->early S
Late S (EdU+) H4 deposition: S-phase
C
Centromeric nucleosome de novo deposition
EYFP-AIDCENP-A
+IAA
New histones incorporation
New histones
Old histones
Ac
H4K5Ac
H4K5Ac CUT&RUN-qPCR at centromeres
NT
IAA
Fold enrichment at alpha-sat compared to IgG & normalized to Alu
1.5
1.0
0.5
0.0
G1-arrested
S-arrested
early S
mid S
late S
G2-arrested
CENP-A depletion at thymidine T = 0
D
Metaphase synchro Colcemid (5h)
5hIAA
M
G2
S
G1
Centromeric CO-FISH
5hIAA
Mitosis
G1
G2
S
Cell synchronization & CENP-A depletion strategy for cen-CO-FISH experiment (fig. 1I)
G2/M synchro RO-3306 (24h)
G1/S synchro Palbociclib (24h)
7hIAA
14hIAA
E
Colcemid
Cdk1i
Palbociclib
A/S
2n
4n
F
From thymidine release
**
*
**
Aberrant Centromeres (% of Total)
100
50
0
NT
IAA
NT
IAA
NT
IAA
Exp1
Exp2
Exp3

Figure S1

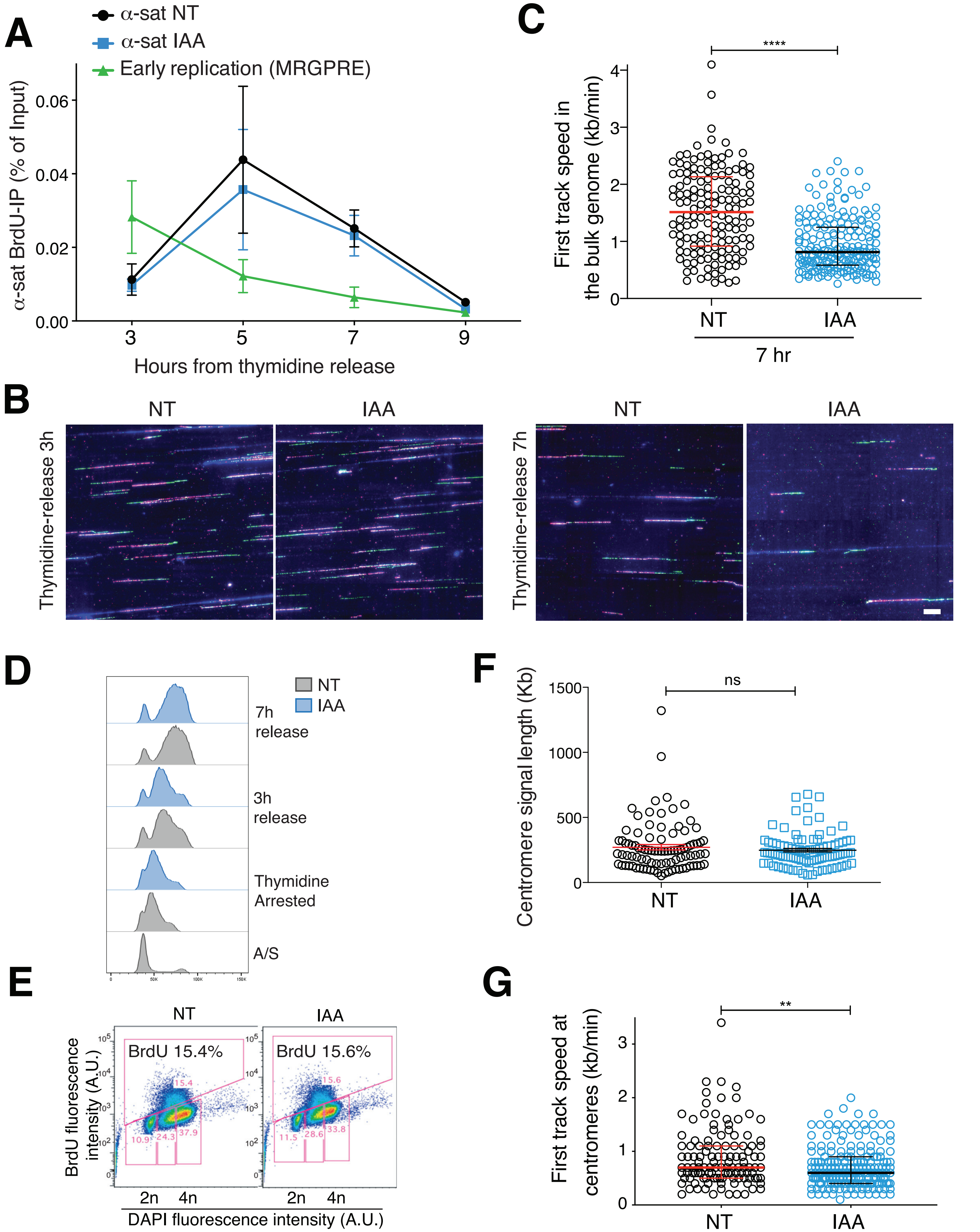
A
α-sat NT
α-sat IAA
Early replication (MRGPRE)
α-sat BrdU-IP (% of Input)
0.06
0.04
0.02
0.00
3
5
7
9
Hours from thymidine release
B
NT
IAA
Thymidine-release 3h
NT
IAA
Thymidine-release 7h
C
****
First track speed in the bulk genome (kb/min)
4
3
2
1
0
NT
IAA
7 hr
D
NT
IAA
7h release
3h release
Thymidine Arrested
A/S
E
NT
IAA
BrdU 15.4%
BrdU 15.6%
15.4
10.9
24.3
37.9
15.6
11.5
28.6
33.8
BrdU fluorescence intensity (A.U.)
2n
4n
2n
4n
DAPI fluorescence intensity (A.U.)
F
ns
Centromere signal length (Kb)
1500
1000
500
0
NT
IAA
G
**
First track speed at centromeres (kb/min)
3
2
1
0
NT
IAA

### Figure S2

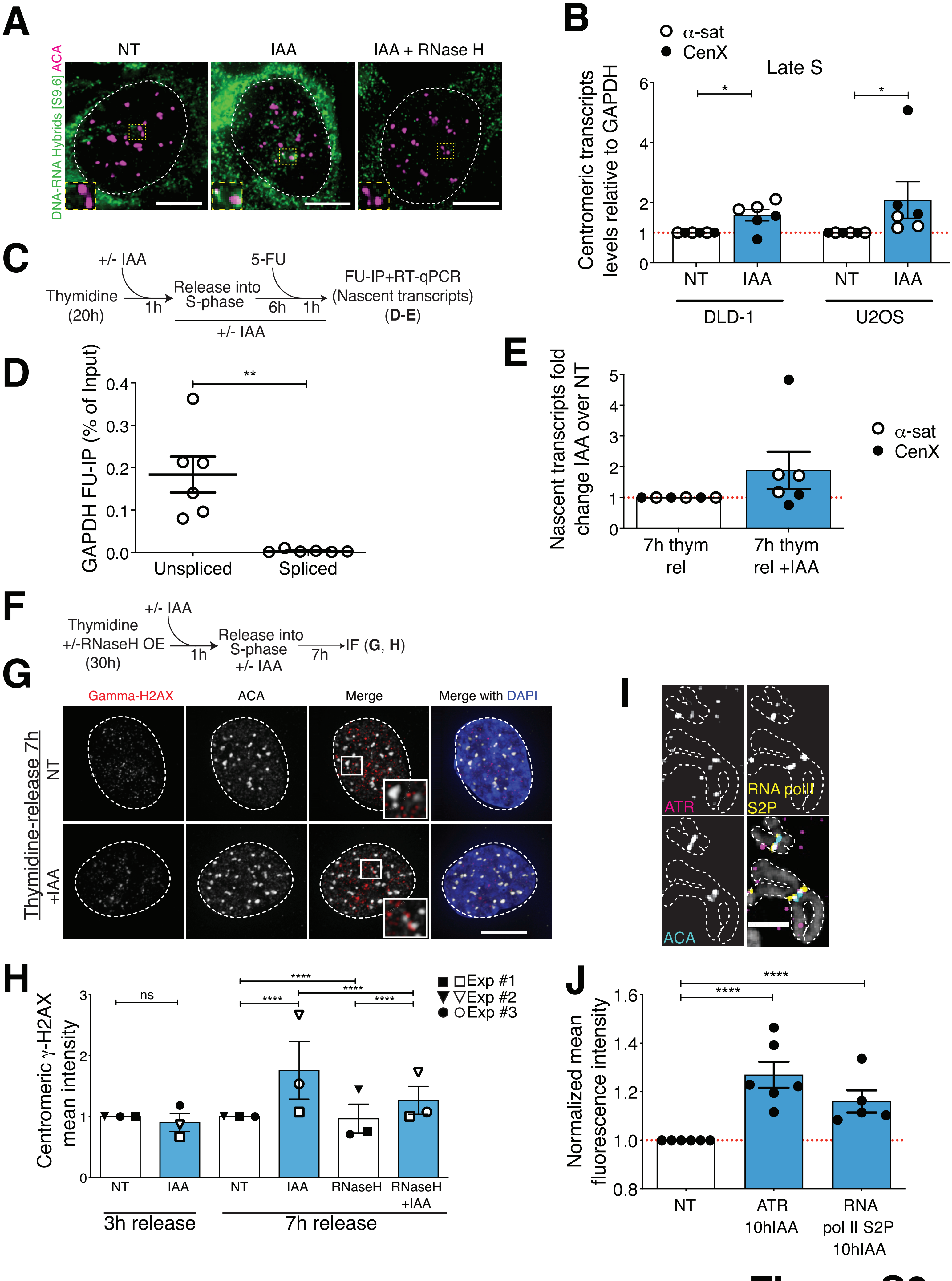

A
NT
IAA
IAA + RNase H
DNA-RNA Hybrids [S9.6] ACA
B
α-sat
CenX
Late S
Centromeric transcripts levels relative to GAPDH
NT
IAA
NT
IAA
DLD-1
U2OS
C
+/- IAA
Thymidine (20h)
1h
Release into S-phase
5-FU
6h
1h
FU-IP+RT-qPCR (Nascent transcripts) (D-E)
+/- IAA
D
GAPDH FU-IP (% of Input)
Unspliced
Spliced
E
Nascent transcripts fold change IAA over NT
7h thym rel
7h thym rel +IAA
α-sat
CenX
F
+/- IAA
Thymidine +/-RNaseH OE (30h)
1h
Release into S-phase +/- IAA
7h
IF (G, H)
G
Gamma-H2AX
ACA
Merge
Merge with DAPI
Thymidine-release 7h
NT
+IAA
H
Centromeric γ-H2AX mean intensity
Exp #1
Exp #2
Exp #3
ns
NT
IAA
NT
IAA
RNaseH
RNaseH +IAA
3h release
7h release
I
ATR
RNA polII S2P
ACA
J
Normalized mean fluorescence intensity
NT
ATR 10hIAA
RNA pol II S2P 10hIAA


### Figure S3

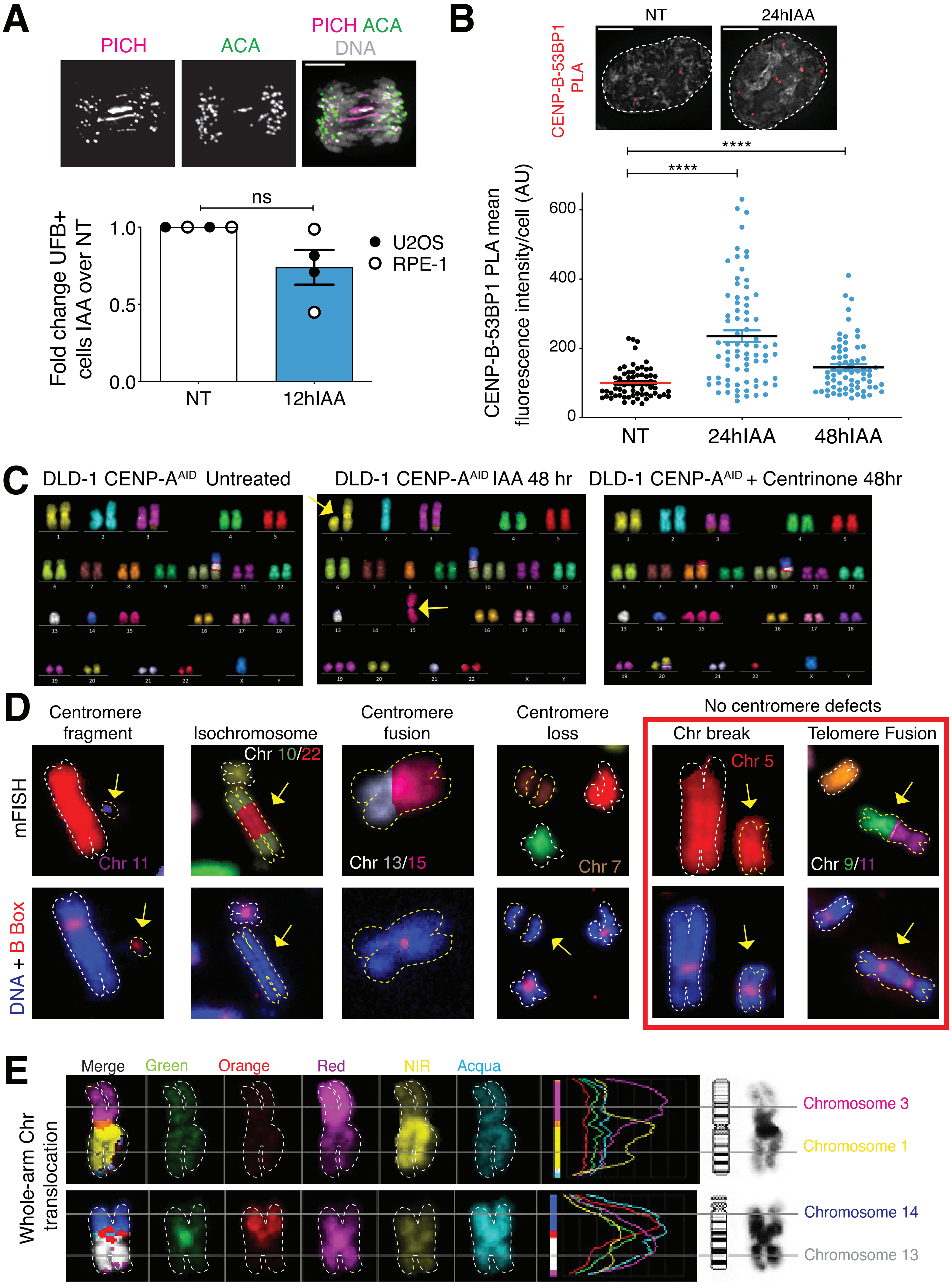

A
PICH
ACA
PICH ACA
DNA
Fold change UFB+ cells IAA over NT
ns
1.0
0.5
0.0
NT
12hIAA
U2OS
RPE-1
B
CENP-B-53BP1 PLA
NT
24hIAA
****
****
CENP-B-53BP1 PLA mean fluorescence intensity/cell (AU)
600
400
200
0
NT
24hIAA
48hIAA
C
DLD-1 CENP-A^AID Untreated
DLD-1 CENP-A^AID IAA 48 hr
DLD-1 CENP-A^AID + Centrinone 48hr
D
Centromere fragment
Isochromosome
Centromere fusion
Centromere loss
No centromere defects
Chr break
Telomere Fusion
mFISH
DNA + B Box
Chr 11
Chr 10/22
Chr 13/15
Chr 7
Chr 5
Chr 9/11
E
Merge
Green
Orange
Red
NIR
Acqua
Whole-arm Chr translocation
Chromosome 3
Chromosome 1
Chromosome 14
Chromosome 13


### Figure S4
